# Chromothripsis as an on-target consequence of CRISPR-Cas9 genome editing

**DOI:** 10.1101/2020.07.13.200998

**Authors:** Mitchell L. Leibowitz, Stamatis Papathanasiou, Phillip A. Doerfler, Logan J. Blaine, Yu Yao, Cheng-Zhong Zhang, Mitchell J. Weiss, David Pellman

## Abstract

Genome editing has promising therapeutic potential for genetic diseases and cancer (1, 2). However, the most practicable current approaches rely on the generation of DNA double-strand breaks (DSBs), which can give rise to a poorly characterized spectrum of structural chromosomal abnormalities. Here, we show that a catastrophic mutational process called chromothripsis is a previously unappreciated consequence of CRISPR-Cas9-mediated DSBs. Chromothripsis is extensive chromosome rearrangement restricted to one or a few chromosomes that can cause human congenital disease and cancer (3–6). Using model cell systems and a genome editing protocol similar to ones in clinical trials (7) (NCT03655678, NCT03745287) we show that CRISPR-Cas9-mediated DNA breaks generate abnormal nuclear structures—micronuclei and chromosome bridges—that trigger chromothripsis. Chromothripsis is an on-target toxicity that may be minimized by cell manipulation protocols or screening but cannot be completely avoided in many genome editing applications.

## Introduction

CRISPR-Cas9 is directed to its target-site by a guide RNA (gRNA), creating specific DNA double-strand breaks (DSBs) almost anywhere in the genome (1, 2). Error-prone DNA repair by non-homologous end joining (NHEJ) of Cas9-generated DSBs can create small insertions and deletions, which can be exploited therapeutically by disrupting protein coding or DNA regulatory sequences. A particularly promising application of this approach is for autologous hematopoietic stem cell (HSC) therapy of common β-hemoglobinopathies including sickle cell disease and β-thalassemia. Specifically, NHEJ-mediated disruption of non-coding DNA regions that repress γ-globin gene (HBG1 and HBG2) transcription can induce fetal hemoglobin (HbF, α2γ2) expression in red blood cell progeny to alleviate the symptoms of sickle cell disease or β-thalassemia (7–9). Cas9 can also be used to initiate precise nucleotide substitutions by homology-directed repair (HDR) for correction of monogenic diseases, including reversion of the mutant sickle cell disease codon (1, 2, 10–13). Several promising CRISPR-based strategies that do not involve the generation of DSBs have been described, but these methods are at an earlier stage of development and have not yet been advanced to clinical trials (14–16).

Because of the clinical potential, it is important to understand the toxicities associated with CRISPR-Cas9 genome editing. Much attention has been paid to unintended, “off-target” DSBs (17). Although well-documented, this outcome can be addressed by the design of more specific gRNAs, novel gene editing strategies, and the development of high-specificity Cas-nucleases (17). Less is known about potential detrimental consequences that arise from on-target genome editing nuclease-mediated DSBs. On-target DNA breakage can induce the *TP53* tumor suppressor and potentially create selective pressure for *TP53* loss followed by tumorigenesis (18–21). Additionally, there have been recent reports of local DNA rearrangements and deletions up to several kilobases in length (22–25), as well as regional megabase-scale deletions telomeric to the on-target DSB (26–28). However, the mechanisms leading to these DNA alterations remain poorly defined, in part because of the lack of high-depth whole genome sequencing. Finally, genome editing protocols involving multiple on-target CRISPR-Cas9 DNA breakage events can lead to incorrect DNA end-joining and reciprocal chromosome translocations that can persist at low levels for months in treated patients (29).

Here, using a variety of approaches including the combination of imaging and single-cell genome sequencing (LookSeq) (30, 31), we report that chromothripsis is a previously unrecognized consequence of on-target Cas9-mediated DNA breakage. This occurs because in actively dividing cells, genome editing with Cas9 causes up to a 28-fold increase in the formation of micronuclei and/or chromosome bridges, aberrant nuclear structures that initiate chromothripsis. In addition to causing rare human congenital disease (5, 32), chromothripsis is common in cancer, where it is well established to generate tumor suppressor loss, fusion oncogenes, or oncogene amplification through the formation of circular double minute chromosomes (3, 6, 33–35). Our findings reveal that initial errors from on-target genome editing can be amplified into far more extensive chromosome alterations in subsequent cell cycles via the generation of aberrant nuclear structures.

## Results

Cas9 generates a DSB that cleaves the targeted chromosome into two segments: one with the centromere region (“centric”) and one without (“acentric”). It seemed likely that if the DSB was not repaired prior to cell division, the acentric fragment lacking a functional centromere would missegregate, forming a micronucleus (Fig. 1a) (4, 36, 37).

**Fig. 1.**
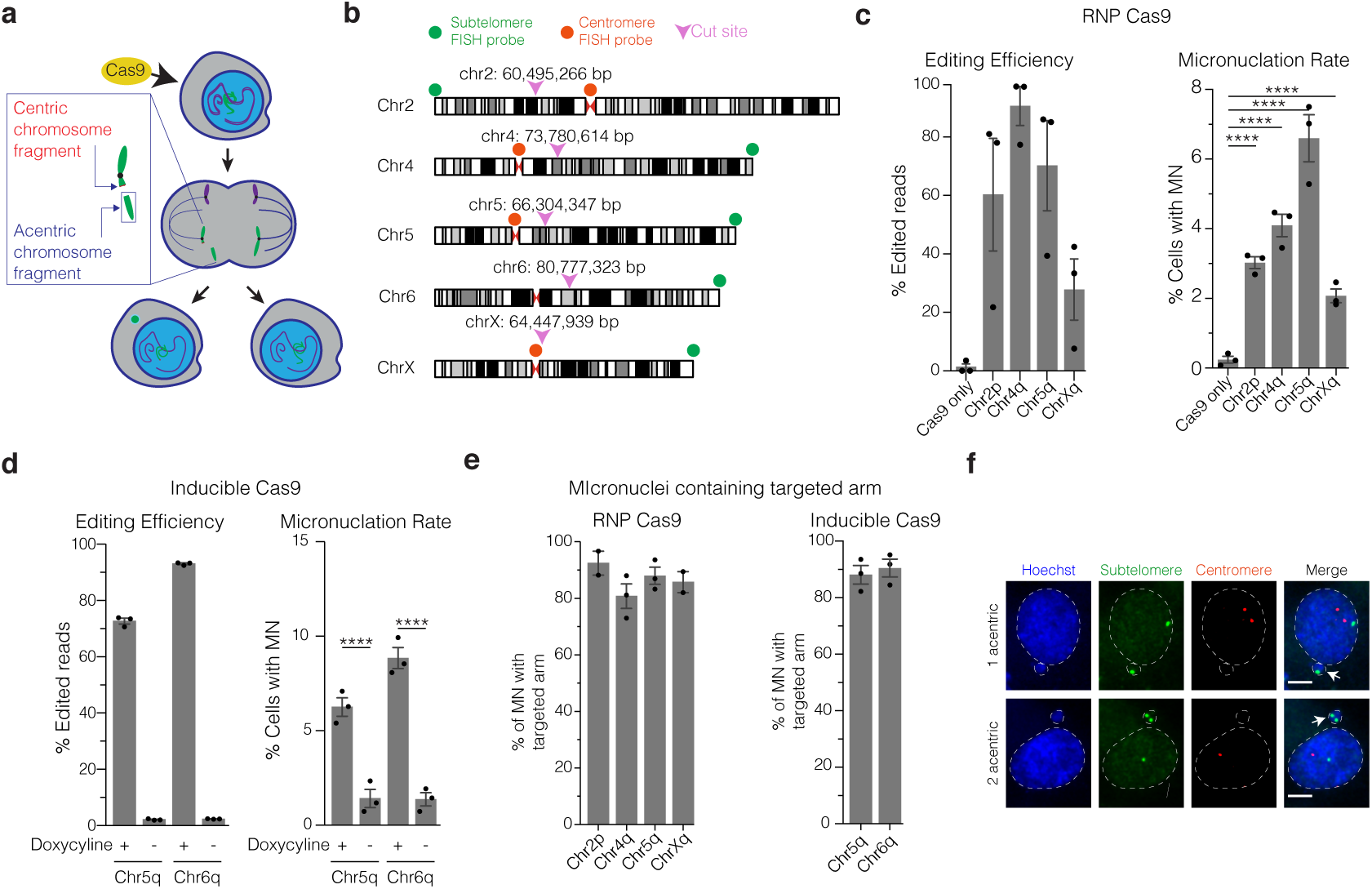
Micronucleation is an on-target consequence of CRISPR-Cas9 genome editing. (a) Schematic of how Cas9 DNA cleavage of a chromosome arm can generate micronuclei. In the shown example DNA cleavage of one sister chromatid occurs in a G2 cell. The centric fragment segregates properly into a daughter nucleus whereas the acentric fragment that cannot be segregated by the spindle is partitioned into a micronucleus. Variations on this outcome include DNA cleavage in G1, cleavage of both sisters in a G2 cell, and cleavage of both homologous chromosomes (not shown). (b) Chromosome locations of the gRNAs and FISH probes used in this study. Magenta arrowheads and numerical coordinates indicate the cut site for specific gRNAs. Green dot: acentric fragment FISH probe locations; red dot: centric fragment FISH probe locations. (c) The frequency of micronucleation after CRISPR-Cas9 RNP transfection. Left, editing efficiency after Cas9/gRNA RNP transfection, 46 h after release of RPE-1 cells from a G1 block. Right, frequency of micronucleation for these RNP transfections. (*n* = 3 experiments with 4007, 4227, 3930, 3325, 3988 cells scored for micronucleation, left to right). Error bars: mean *±* SEM, *P <* 0.0001, two-tailed Fisher’s exact test. (d) As in panel C, but for doxycycline-inducible CRISPR-Cas9 with constitutively expressed gRNA. p53 siRNA treatment was performed prior to doxycycline treatment. (*n* = 3 experiments with 1265, 1261, 1244, 1239 cells scored for micronucleation, left to right). Error bars: mean *±* SEM, *P <* 0.0001, two-tailed Fisher’s exact test. (e) Percentage of MN containing the targeted chromosome arm. Left, RNP transfection (*n* = 2 experiments with 64 and 96 cells scored for chr2p, chrXq, respectively, *n* = 3 experiments with 83 and 116 cells scored for chr4q, chr5q, respectively). Right, RPE-1 cells with inducible-Cas9 and constitutively expressed gRNA (*n* = 3 experiments with 168 cells scored for each). (f) Example images of FISH analysis after Cas9/gRNA RNP transfection (single plane from a confocal imaging stack). Red channel: centric fragment probe; green channel: acentric fragment probe; blue channel: Hoechst stain (DNA); white arrows: micronuclei; dashed white line: outline of Hoechst (DNA) label. Scale bar 5 µm.

We first evaluated this possibility in genetically stable human retinal pigment epithelial cells (hTERT-RPE-1). To estimate the rate of micronucleation in a single cell cycle, we synchronized cells with a serum starvation-block and release protocol followed by transfection with a Cas9/gRNA ribonucleoprotein (RNP) complex shortly before the next cell division (22 hours after release, approximately during S/G2 [Supplementary Fig. S1a]). We used single guide RNAs (gRNAs), each targeting unique genomic sites on four different chromosomes (Fig. 1b and Supplementary Table S1). All gRNAs targeted intergenic sequences, except one that disrupts the erythroid-specific enhancer of the *BCL11A* gene on chromosome 2 (“chr2p”), similar to strategies that are currently in clinical trials for β-thalassemia and sickle cell disease (7, 38) (NCT03655678, NCT03745287). The *BCL11A* gene encodes a transcriptional repressor protein that silences γ-globin expression postnatally in red blood cells. CRISPR-Cas9 cutting at individual target sites induced micronucleation at frequencies of 2.1–6.6%, 8.6-to 27.5-fold higher than controls (hereafter “CRISPR-MN”, [Fig. 1c]). Similar results were obtained using cells that constitutively express the gRNA targeting chr5q and a fifth gRNA targeting chr6q, where Cas9 was conditionally expressed from a thirdgeneration doxycycline-inducible promoter (Fig. 1b,d and Supplementary Fig. S1a,b) (39).

Fluorescence in situ hybridization (FISH) established that 77–92% of CRISPR-MN contained the chromosome arm targeted by the specific gRNAs (Fig. 1e,f and Supplementary Fig. S1c). Most micronuclei contained two copies of the targeted chromosome segment, which could result from either cleavage of both homologous chromosomes in a G1 cell, or from cleavage of both sister chromatids of one homolog in a G2 phase cell (Fig. 1f and Supplementary Fig. S1c). Co-staining with centromere-specific FISH probes confirmed that CRISPR-MN are mostly acentric chromosome fragments (Fig. 1f). Similar results for micronucleus formation and chromosome arm copy number alterations were also obtained in BJ foreskin fibroblasts (Supplementary Fig. S1d,e).

CRISPR-MN exhibited defects in nuclear function observed previously in micronuclei, including spontaneous nuclear envelope rupture (Supplementary Fig. S1f), defective DNA replication (Supplementary Fig. S1g), and the accumulation of DNA damage (Supplementary Fig. S1h-j) (40–47). Therefore, CRISPR-Cas9 genome editing can generate micronuclei containing the acentric fragment of the targeted chromosome, which is then subject to extensive DNA damage.

These findings suggested that chromothripsis might be an unrecognized, on-target consequence of CRISPR-Cas9 genome editing. To directly test this hypothesis, we used “Look-Seq”, a procedure combining long-term livecell imaging with single-cell whole-genome sequencing of the imaged cells (30, 31). CRISPR-MN were generated in daughter cells as above. Because Cas9 genome editing can be limited by p53 induction (18–21), we transiently depleted p53 by siRNA-mediated knockdown prior to inducing CRISPR-MN. Micronucleated daughter cells were allowed to divide, and their progeny (granddaughter cells) were then isolated for single-cell sequencing.

In total, we sequenced 18 granddaughter pairs derived from micronucleated daughter cells (Fig. 2a); we tested three different guides, including the chr2p guide targeting the erythroid-specific *BCL11A* enhancer (7, 38). The micronucleated acentric chromosome segments exhibited several patterns of copy number alterations, which can be explained as follows. Cas9 can cleave one or both homologous chromosomes and one or both sister chromatids (the gRNAs in this study were not designed to be allele-specific); acentric fragments can be distributed in any combination to granddaughter cells; and/or the micronuclear DNA can be severely underreplicated (in most cases, DNA replication in micronuclei is highly inefficient) (4, 44, 46). As shown in Fig. 2a, we observed examples consistent with each of the above scenarios, as well as additional factors discussed below.

**Fig. 2.**
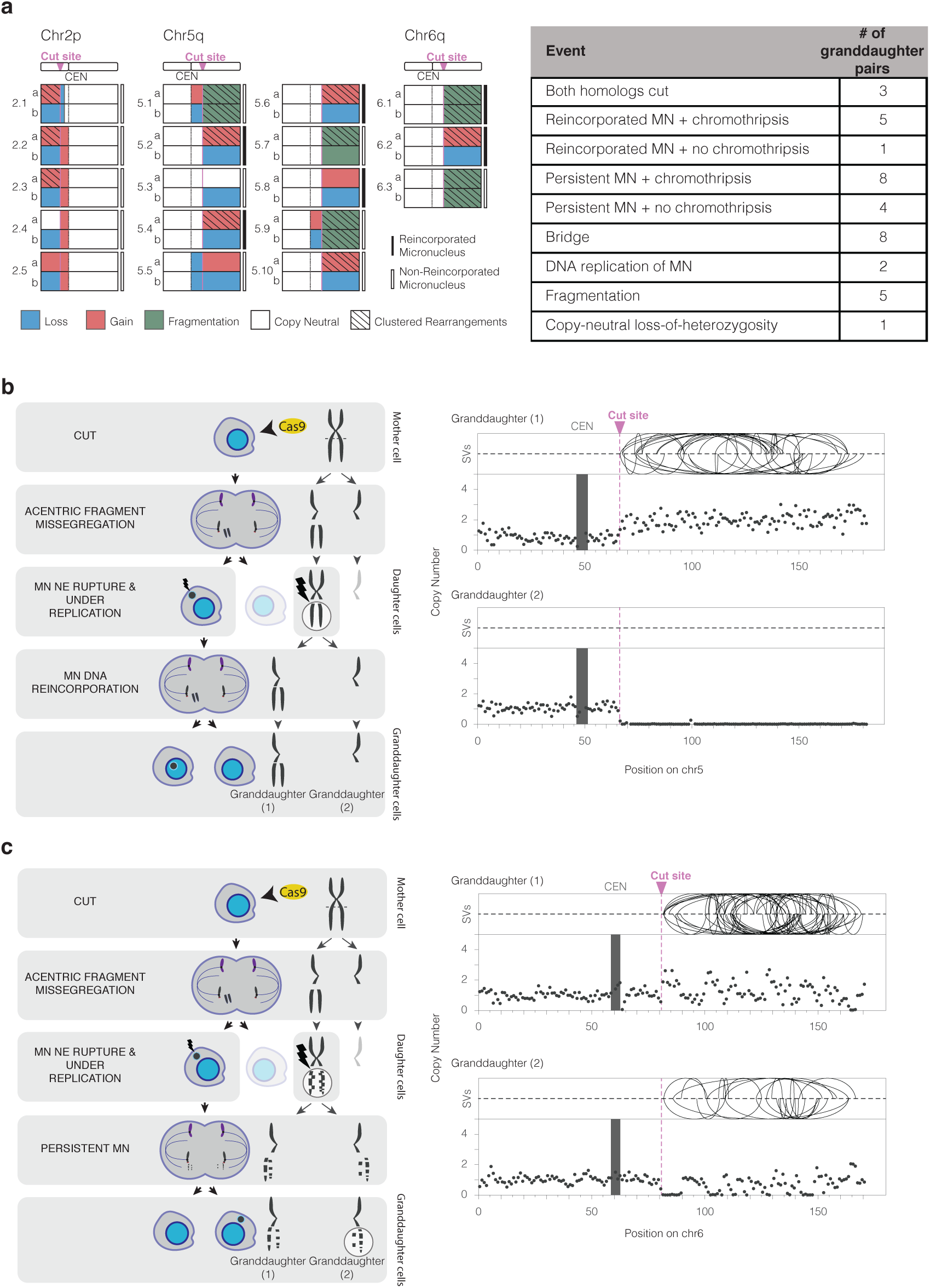
CRISPR-Cas9 genome editing can cause chromothripsis. (a) Summary of genomic outcomes after the division of 18 micronucleated cells. Left, schematic summary providing a more detailed view of each the 18 daughter cell pairs. Number to the left of the schematics is an ID: first number is the targeted chromosome; second number is a sample identifier for that chromosome. Right, summary table. In 3/18 samples there was CRISPR-Cas9 cleavage of both homologs. Reincorporation of the MN DNA was inferred from the absence of detectable GFP-H2B signal in either granddaughter. Bridges were inferred from the DNA sequence analysis. Replication of the MN or copy-number neutral LOH was inferred from haplotype-resolved DNA copy number. Fragmentation is evident from reciprocal changes in the DNA copy number along the chromosome arm when comparing the two granddaughters. Note that a given sample can have more than one genomic outcome. (b) Chromothripsis after a micronucleus is reincorporated into a granddaughter cell. Left, cartoon depicting the cellular events leading to the genomic outcomes for CRISPR-Cas9 sample 5.6 (Supplementary Fig. S3). Cells are on the left and chromosomes are depicted on the right. In the first generation, both sisters from one homolog were cleaved in a G2 cell (horizontal dashed line) that divides to generate a micronucleated daughter (left) and a non-micronucleated daughter (right, faded cell not subsequently followed). DNA in the micronucleus is poorly replicated. In the second cell division, the micronuclear chromosome is reincorporated into a granddaughter cell’s primary nucleus. Lightning bolt: DNA damage. Right, plots showing structural variants (SVs) and DNA copy number for haplotype of the cleaved chromosome. Top, intrachromosomal SVs (> 1 Mb) are show by the curved lines. Bottom: copy number plot (1 Mb bins). CEN: centromere. (c) Chromothripsis after the bulk of a micronuclear chromosome fails to be reincorporated into a granddaughter cell primary nucleus for sample 6.3 (Supplementary Fig. S3). Cartoon (left) and SV and copy number plots (right) as in (b). In this example, the two arms from cleaved sister chromatids are fragmented, generating chromothripsis in both daughters. *TP53*

Haplotype-specific copy number analysis showed that in 15 of 18 granddaughter pairs, the acentric arm from one homologous chromosome was missegregated, whereas both homologs were missegregated in the remaining 3 pairs (Fig. 2a). In one notable example, missegregation of both homologs to opposite daughters led to copy-neutral loss-of-heterozygosity (LOH; i.e., uniparental disomy for the acentric fragment in both granddaughter cells, sample 5.1 in Supplementary Fig. S2, data in Fig. 3b). Copy-neutral LOH is common in cancer and can result in tumor suppressor inactivation (48). Our findings therefore provide a clear mechanistic explanation for similar patterns that had been noted, but not explained, after CRISPR-Cas9 genome editing (28, 49, 50). In summary, on-target Cas9 genome editing can generate micronuclei, which in turn can induce large-scale DNA copy number alterations as well as copy-number neutral LOH.

**Fig. 3.**
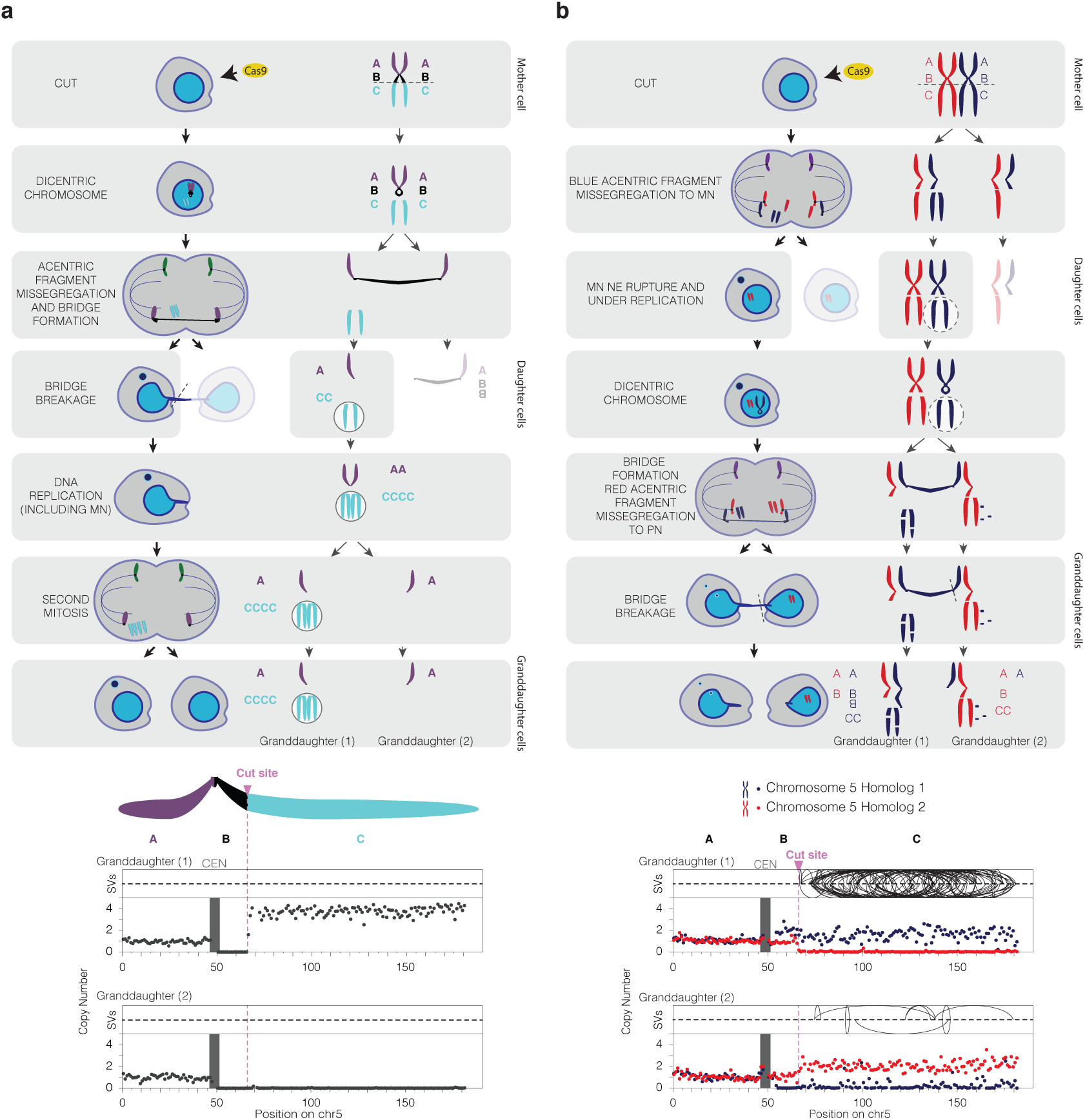
CRISPR Cas9-genome editing induces chromosome bridge formation, adding to the genome complexity from micronuclei. (a) Evidence for genome editing-induced chromosome bridge in sample 5.5 (Supplementary Fig. S4). Scheme as in Fig. 2. CRISPR-Cas9 cut site is indicated by the dashed line and relevant segments of chr5 are indicated by letters A-C. In the first division, the DNA break on sister chromatids results in the formation of a micronucleus with the acentric portions of chr5 (segment C). At the same time, the sister centric fragments (AB) fuse, generating a dicentric bridge concomitantly with the formation of the micronucleus. Asymmetric breakage of the bridge leads to the loss of the “B” segment from the bridge chromosome in the micronucleated daughter. Faded cell inferred to contain two copies of the B segment was not followed further. DNA copy number analysis indicated that in this example the chromosome fragments in the micronucleus underwent DNA replication. This region showed no detectable rearrangements. Note that the acentric fragments of chr5 were not reincorporated into a daughter primary nucleus in the second division. A (purple): p-arm; B (black): centromere to cut site, inferred to reside in the bridge; C (teal): cut site to the telomere. Bottom: Copy number and rearrangement plots of cells from above, as in Fig. 2. (b) Bridge formation, micronucleation, chromosome fragmentation and chromothripsis from CRISPR-Cas9 genome editing in sample 5.1 (Supplementary Fig. S3). In this sample both homologs were cleaved. The acentric arm of homolog 1 (blue allele) missegregates into a micronucleus in the first generation. The centric fragments of homolog 1 fuse, resulting in a dicentric bridge in the second cell division, as in (A). After the second cell division, the cell that inherited the acentric fragment of homolog 2 (red) was found to have few SVs or copy number alterations, suggesting it was partitioned into the primary nucleus as indicated in the scheme. By contrast, the acentric segments of homolog 1 were fragmented. Bottom: copy number and rearrangement plots of cells shown above, as in Fig. 2.

Chromothripsis is extensive chromosomal rearrangements that are clustered on one or a few chromosomes or chromosome arms, and are commonly accompanied by oscillations between two or three DNA copy number levels (4, 6, 51). We identified the characteristic clustering of rearrangements on the acentric segment of the Cas9-targeted chromosome arm in 13 of 18 granddaughter pairs sequenced (Fig. 2a,b and Supplementary Figs. S2-S3, *P* ≤ 10^*–*7^, onesided Poisson test). The most striking example was a targeted chr6q arm in which we detected 646 intrachromosomal breakpoints distributed between the two granddaughter cells (Supplementary Figs. S2-S3, sample 6.1).

Haplotype copy number analysis also demonstrated fragmentation of the targeted acentric chromosome fragment. If a chromosome from a micronucleus is fragmented, and the fragments are distributed randomly between the granddaughter cells, the granddaughter cells will display a mirror image DNA copy number pattern that oscillates between two levels (31). In the simplest case, one homolog is replicated and segregated normally. However, the fragmentation of the other homolog can generate oscillations between zero copies and one copy in each daughter. The regions with one copy of the fragmented homolog will retain heterozygosity, leading to islands of heterozygosity interspersed within regions of LOH, one criterion for chromothripsis (51). In five of 18 pairs we observed fragmentation based on haplotype copy number analysis (Fig. 2a and samples 5.1, 5.7, 5.9, 6.1, 6.3 in Supplementary Fig. S2 and data in Fig. 3b). In eight of 18 pairs, there were clustered rearrangements on the targeted arm without detectable copy number oscillations, producing copy-neutral chromothripsis. In the copy-neutral cases, the acentric segment was fragmented but most fragments were inherited by only one granddaughter. Copy-neutral chromothripsis is frequently observed in human congenital diseases, an observation that is likely explained by the strong selection against gene copy number imbalance during human development (32).

Micronuclei that spontaneously lose their nuclear envelope integrity demonstrate defects in nuclear functions, such as transcription and DNA replication (41, 44). Accordingly, we and others previously hypothesized that the DNA ligation required to generate chromothripsis would only occur after mitosis and upon reincorporation of the micronuclear chromosome into a nucleus with functional DNA end-joining (31, 45, 52). However, in many cases, micronuclear chromosomes fail to reincorporate into a primary nucleus and are again partitioned into micronuclei (37, 40). Furthermore, micronuclei that lack kinetochores, like CRISPR-MN, are rarely reincorporated (37, 52).

We tested whether bulk chromosome reincorporation is required to generate chromothripsis by sequencing granddaughter cells with micronuclear chromosomes present in the cytoplasm. Surprisingly, of the 12 CRISPR-generated granddaughter pairs with a persistent micronucleus, eight showed chromothripsis involving the targeted chromosome arm (Fig. 2a,c, Supplementary Fig. S2 and Supplementary Videos 1,2). Chromothripsis in these samples could either be due to endjoining of chromosome fragments in the cytoplasm, aberrant mitotic DNA synthesis (30, 53), or ligation of a subset of chromosome fragments that might have been incorporated into the granddaughter nucleus after the division of a micronucleated cell. It was recently reported that spontaneously arising micronuclei in mouse embryonic cells often fail to be reincorporated, which was hypothesized to reflect a mechanism to prevent chromothripsis during embryo development (52). However, our data establish that chromothripsis can occur even without visible reincorporation of micronuclear chromosomes.

These results provide an important validation of our previous single-cell analysis showing that micronuclei can cause chromothripsis. In this prior work, we used random mitotic errors to generate micronuclei and then inferred the identity of the micronuclear chromosome based on it being the only underreplicated chromosome (31). The current results, in which the identity of the micronuclear chromosome is known a priori, confirm that the micronuclear chromosome is the one that undergoes chromothripsis. Moreover, in 11 of the 13 samples with chromothripsis, haplotype-resolved DNA copy number analysis demonstrated that the micronuclear chromosome showed little detectable replication, again confirming our prior work (31). Together with other recent work examining clonal cell populations after the induction of micronuclei, it is now clear that these structures generate chromothripsis at remarkably high rates (34, 43, 45).

In addition to the formation of micronuclei, Cas9-generated DNA breaks can lead to dicentric chromosome bridge formation due to ligation of the centric fragments of Cas9-cleaved sister chromatids (30, 54). We recently delineated a series of mechanistic steps through which chromosome bridges, like micronuclei, induce chromothripsis (30).

In eight out of 18 pairs derived from CRISPR-MN cells, we identified signatures of bridge formation, which added complexity to the copy number patterns resulting from the missegregation of acentric fragments as described above. All of these samples involve two cell divisions during which bridges could form (Supplementary Fig. S4): the first division, when the micronucleus is generated; and the second division, when the micronucleated daughter cell divides. If the bridge forms and breaks in the first cell division, the granddaughters will exhibit shared segmental gains or losses on the centromeric side of the Cas9 cut, as seen in six of 18 granddaughter pairs (Fig. 3a and Supplementary Fig. S4a). If the bridge forms and breaks in the second cell division, the cells will display reciprocal gain and loss of DNA sequence on the centromeric side of the cut site, as observed in two of 18 granddaughter pairs (Fig. 3b and Supplementary Fig. S4b). Note that the megabase-scale copy number loss on the centromeric side of the cut, which we attribute to bridge breakage, cannot be explained by DNA resection from the cut site because resection is generally limited to several kilobases (55). Moreover, resection cannot explain copy number gains on the centromeric side of the breaks. Instead, segmental gains are a sequence signature of the chromosome breakagefusion-bridge cycle, a common mutational process in cancer that generates gene amplification (30, 56–59). Finally, the chromosome that was inferred to form a bridge shared the same haplotype as the micronuclear chromosome. This is in agreement with the expectation that dicentric bridges and acentric micronuclei can arise simultaneously from the same Cas9 cut.

Support for chromosome bridge formation also came from fluorescence imaging, which showed that 10.7% of cell divisions that formed micronuclei after CRISPR-Cas9 cutting also formed visibly detectable chromosome bridges (Supplementary Fig. S4c). Micronucleation physically separates the centric and acentric fragments, preventing the acentric fragment from being used as a ligation partner for the centric fragment of the broken chromosome. We reasoned that the presence of a micronucleus would be expected to create a bias for ligation of the centric fragments of the broken chromosome, leading to elevated rates of dicentric bridge formation in the granddaughters. Consistent with this, the frequency of bridge formation was higher still after the division of micronucleated cells (27.2% of divisions [Supplementary Fig. S4c]). Thus, CRISPR-Cas9 genome editing is accompanied by chromosome bridge formation in addition to micronucleation, both of which can trigger ongoing cycles of genome instability.

Lastly, we tested the applicability of our findings to a protocol being investigated for the treatment of β-hemoglobinopathies (7–9). We electroporated human CD34+ hematopoietic stem and progenitor cells (HSPCs) with a Cas9/gRNA RNP complex targeting the erythroid-specific *BCL11A* enhancer on chromosome 2p (7, 38). We confirmed successful editing and concomitant increase of HbF expression in erythroid progeny (Fig. 4a,b). Similar to our observations in cell lines, the frequency of micronucleation increased 16-fold by 24 hours after RNP transfection into CD34+ HSPCs (Fig. 4c). Using FISH probes surrounding the Cas9 cut, we found that over 80% of cells containing micronuclei exhibited copy number alterations affecting the acentric fragment of the targeted chromosome (Fig. 4d,e). Moreover, 7.3% of cells without micronuclei exhibited abnormal numbers of this chromosome arm, indicating cutting that is followed by missegregation of the acentric fragment to the primary nucleus (Fig. 4d,e). Some of these *TP53*intact cells were capable of entering mitosis with an unrepaired DNA break, as 3.25% of cells analyzed had breaks in chr2p detected by spectral karyotyping (SKY) 24 h after Cas9 treatment (Fig. 4f,g). We also detected high-level phosphorylation of histone H2AX in 12.9% of micronuclei (Fig. 4h,i), indicating extensive DNA damage. Although it is not feasible to apply Look-Seq single-cell sequencing to non-adherent cells such as HSPCs, our results establish that these cells acquire hallmark cytological features associated with chromothripsis following CRISPR-Cas9 genome editing.

**Fig. 4.**
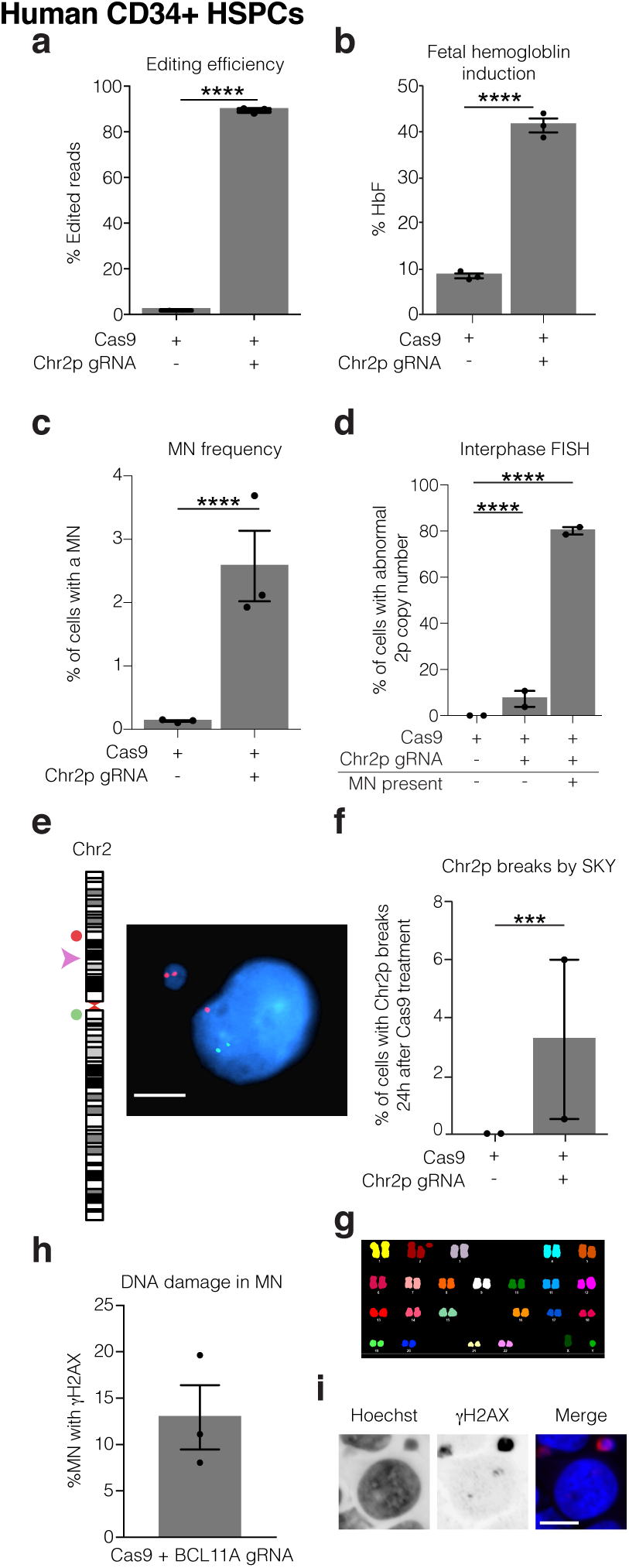
Hallmark cytological features of chromothripsis after a genome editing protocol for the treatment of sickle cell disease. Human CD34+ HSPCs were electroporated with Cas9/gRNA RNP targeting the erythroid-specific enhancer of *BCL11A*. Microscopic analysis of micronucleation was performed 24 h post electroporation. (a) Editing efficiency of *BCL11A* determined from amplicon sequencing. Error bars: mean *±* SEM, *P <* 0.0001, two-tailed unpaired *t* -test). (b) Fetal hemoglobin (HbF) levels were measured by HPLC in erythroid-differentiated CD34+ HSPCs as a functional readout of successful editing of *BCL11A* 10 days after RNP electroporation. Error bars: mean *±* SEM, test *P <* 0.0001, two-tailed unpaired *t* -test). (c) Percent of cells with a micronucleus (*n* = 3 experiments with 7827 and 6480 cells counted, left to right). Error bars: mean *±* SEM, *P <* 0.0001, two-tailed Fisher’s exact test. (d) Percent of cells with aberrant 2p copy number assayed by FISH (*n* = 2 experiments with 1957, 1926, 74, cells counted, left to right). Error bars: mean *±* SEM, *P <* 0.0001, two-tailed Fisher’s exact test. (e) Representative FISH image for data in (d). Cut site is represented by a pink arrow; DNA is blue; telomere proximal probe is red; centromere proximal probe is green. Shown is a micronucleated cell with 3 copies of the cut arm, two of which are in the micronucleus. Scale bar 5 µm. (f) Chr2p breaks present 24 hours after electroporation in metaphase visualized by SKY (*n* = 2 experiments, 400 spreads per condition). Error bars: mean *±* SEM, *P <* 0.0001, two-tailed Fisher’s exact test. (g) Sample SKY image from (F). (h) Percent of CD34+ CRISPR-MN with extensive DNA damage covering the DNA present in the micronucleus by γH2AX-labeling (*n* = 3 experiments, 135 micronuclei scored) (i) Representative image of data in (h). Scale bar 5 µm.

## Discussion

Here we demonstrate that on-target CRISPR-Cas9 genome editing can induce the formation of micronuclei and chromosome bridges in dividing cells, leading to copy number alterations of large chromosomal segments and chromothripsis. These findings may, in part, explain the mechanisms underlying recently observed large chromosomal deletions or loss of heterozygosity surrounding on-target DSBs following genome editing in embryos (26, 50, 60–62). Moreover, they raise a new potential concern for therapeutic genome editing strategies that require DSB formation, because chromothripsis can drive the rapid acquisition of multiple cancer-causing mutations simultaneously (4, 6, 33).

In quantitative terms, the risk of chromothripsis after therapeutic genome editing in human subjects remains unclear. Most chromothripsis events should compromise cell fitness, leading to senescence or cell death. To date, malignant transformation or clonal cell expansion following genome editing has not been observed in animal studies, including non-human primate models (63–65), nor in human subjects who have participated in clinical trials (29). Although the survival of cells with chromothripsis is likely promoted by *TP53* loss, it is clear that chromothripsis can occur and persist in *TP53* proficient cells (32, 33, 35). Patients with congenital disease caused by chromothripsis do not have loss of *TP53* (32), and it has even been reported that chromothripsis can revert a dominant genetic disease, enabling hematopoietic stem cell proliferation (66). Similarly, clonal expansion of malignant cells with chromothripsis is common. However, even across human cancers, the incidence of chromothripsis is only enriched 1.5-fold in *TP53* mutant tumors (33).

Our results have a number of practical implications. Efficient Cas9-mediated HDR requires cells to be actively dividing whereas NHEJ does not. Therefore, therapeutic genome editing via NHEJ in non-dividing cells, such as retinal photoreceptors (67), should not produce micronuclei. Conversely, efforts to specifically edit dividing cells in order to enhance HDR rates, for example, by using a modified Cas9 with reduced activity in non-dividing cells (68), may enhance micronucleation and its downstream consequences, including chromothripsis. Accordingly, we suggest that for NHEJ-dependent strategies for therapeutic editing of HSCs, it may be beneficial to maintain HSC quiescence. Some CD34+ HSPC editing protocols appear to favor quiescent or G1 HSCs, whereas other protocols cause a higher frequency of editing in cycling or G2 HSCs (7, 11, 65, 68–70). Additionally, we suggest that for NHEJ applications, fusion of Cas9 to a G1-specific Cdt1 segment could be employed to restrict editing to G1 cells (68, 71), thereby minimizing the probability of micronucleus formation, chromosome segment missegregation during mitosis, and chromothripsis. Screening for micronucleation and/or chromothripsis in clinical protocols should also be considered. Such screening is expected to become more feasible as high-throughput and low-cost methods for single-cell genome sequencing are developed (72). Finally, our study further motivates the development of genome editing strategies that do not generate double-stranded DNA breaks (1, 14–16, 73), which should in principle minimize the potential for inducing chromothripsis.

## Supporting information

Supplementary Video 1_reincorporated.mov

Supplementary Video 2_nonreincorporated.mov

## ACKNOWLEDGEMENTS

We are grateful to R. Jaenisch, S. Markoulaki, A. Spektor and members of the Pellman and Weiss laboratories for discussions, and I. Cheeseman for the doxycycline-inducible Cas9 RPE-1 cell line. This material is based upon work supported by the National Science Foundation Graduate Research Fellowship under Grant No. DGE1144152 (M.L.L.), National Cancer Institute (NCI) career transition award K22CA216319 (C.-Z.Z.), Howard Hughes Medical Institute (D.P.), NIH grant R01 CA213404 (D.P.), F32 DK118822 (P.A.D), NIH grant P01 HL053749 (M.J.W.), the Assisi Foundation (M.J.W.), the Doris Duke Charitable Foundation (M.J.W.), and St. Jude/ALSAC. The St. Jude Cytogenetic Shared Resource Laboratory is supported by NIH grant P30 CA21765 and by St. Jude/ALSAC. We would like to thank the members of the St. Jude Children’s Research Hospital Center for Advanced Genome Engineering and Cytogenetics core facilities.

## AUTHOR CONTRIBUTIONS

M.L.L., S.P. and D.P. conceived the project, M.L.L., S.P., D.P., and P.A.D. designed the experiments, M.L.L. and S.P. performed the experiments except the human blood cells experiments that were carried out by P.A.D. and Y.Y.; M.L.L., S.P., L.J.B. and C-Z.Z. analyzed data, C-Z.Z. and L.J.B. developed computational methodology, M.L.L., S.P. and D.P. wrote the manuscript, all authors discussed the results and commented on the manuscript, M.J.W. supervised the human blood cells experiments, D.P. supervised the study.

## COMPETING INTERESTS

M.J.W. is a consultant for Rubius Inc., Cellarity Inc., Beam Therapeutics, and Esperion; none of the consulting work is relevant to the current project. C.-Z. Z. is a scientific adviser for Pillar BioSciences. All other authors declare no competing interests.

## DATA AVAILABILITY

All datasets and cell lines generated and/or analyzed in the current study are available from the corresponding author upon reasonable request. These include original images, videos and/or analyses for Figs. 1c-f, and 4a-i and Supplementary Figs. S1b-j, S3, and S4c. CD34+ HSPC-derived FISH and SKY images and analyses (Fig. 4d-g) were generated by The St. Jude Cytogenetic Shared Resource Laboratory and derived data supporting the findings in Fig. 4d-g are available from the corresponding author upon request. All Look-Seq videos are available upon request. All sequencing data will be made available in the Sequencing Read Archive (SRA) upon acceptance. All SV calls are also available upon request.

## CODE AVAILABILITY

Bioinformatics pipeline code used for processing of sequencing data is available from the corresponding author upon reasonable request, and is expounded on in Umbreit et al., 2020.

## Methods

### Cell culture and generation of cell lines

Cells were cultured at 37 ºC in 5% CO2. Telomerase-immortalized RPE-1 retinal pigment epithelium and BJ foreskin fibroblasts from ATCC were grown in Delbucco’s Modified Eagle Medium/F12 (1:1) (Gibco) with 10% FBS, 100 IU/ml penicillin and 100 µg/ml streptomycin. RPE-1 cells expressing Cas9 under a doxycycline-inducible promoter (gift from I. Cheeseman (39)) were grown using tetracycline-free FBS (X&Y Cell Culture). Mobilized peripheral blood CD34+ cells were obtained from three de-identified healthy donors (Key Biologics, Lifeblood) and enriched by immunomagnetic bead selection using an AutoMACS instrument (Miltenyi Biotec). Cryopreserved CD34+ cells were thawed and pre-stimulated for 48 h in StemSpan SFEM (StemCell Technologies) supplemented with 100 ng/mL SCF, FLT3L, and TPO (R&D Systems). CD34+ cells were maintained in complete SFEM post-electroporation for 1-5 days or subject to erythroid differentiation. Erythroid differentiation was induced using a two-phase protocol. Phase 1 (days 0-5): IMDM (Thermo) supplemented with 20% FBS, 1% penicillin/streptomycin, 20 ng/mL SCF, 1 ng/mL IL3 (R&D Systems), and 2 U/mL EPO (Amgen). Phase 2 (days 5-10): IMDM supplemented with 20% FBS, 1% penicillin/streptomycin, 2 U/mL EPO, and 0.2 mg/mL holotransferrin (Millipore Sigma).

RPE-1 cells expressing H2B-eGFP, TDRFP-NLS, and eGFP-BAF were created by transduction of lentivirus or retrovirus vectors containing the genes of interest. Virus was generated by transfection of HEK293FT cells with appropriate packaging plasmids (Lentivirus: pMD2.G and psPAX2; Retrovirus: pUMVC and pVSV-G) with Lipofectamine 2000 (Life Technologies) according to the manufacturer’s instructions. Cells were virally transduced for 16 h in the presence of 10 µg/ml polybrene and populations of transduced cells were selected by fluorescence activated cell sorting 7 days later.

### Cas9 RNP transfection in immortalized cell lines

sgRNAs were synthesized with the Trueguide Synthetic gRNA platform (Thermo Fisher Scientific) as chemically modified custom oligos, where the final 3 bases on both the 5’ and 3’ end of the sgRNA are 2’ -O-Methyl bases and the linkages between them are phosphorothioates, in order to increase editing efficiency and protect from nuclease degradation. Their sequences are listed in Supplementary Table S1.

RNP complexes were prepared following a modified version of the suggested manufacturer’s protocol. Briefly, gRNA/Cas9 complexes were formed by incubating 250 ng of the gRNA with 1 µg of purified Cas9 protein (TrueCut Cas9 Protein v2, Invitrogen) in OptiMEM (Invitrogen). Cells were seeded on 12-well dishes, #1.5 glass coverslips (fixed imaging experiments), or 35-mm gridded ibiTreat dishes (ibidi) (Look-Seq), were synchronized by serum starvation in 0.1% FBS-containing media for 24 h and subjected to Cas9 RNP transfection 22 h upon release from the block. Transfection of ribonucleoprotein complexes was performed using Lipofectamine CRISPRMAX Reagent (Invitrogen), following the manufacturer’s protocol. Cells were fixed 46 h after release from block to measure the percentage of cells with micronuclei and ∼35 h after release for the FISH experiments.

### Editing of CD34+ HSPCs

Purified recombinant Cas9 protein was obtained from Berkeley Macrolabs. Chemically modified single guide RNAs (sgRNA) were synthesized by Synthego with 2’ -O-methyl 3’ -phosphorothioate modifications between the 3 terminal nucleotides at both the 5’ and 3’ ends. Ribonucleoprotein complexes (RNPs) were formed by incubating Cas9 (32 pmol/100,000 cells) with sgRNAs at a 1:2 molar ratio. CD34+ cells were washed in PBS, resuspended in the manufacturer provided buffer for primary cells, mixed with RNPs, and electroporated using program 24 of a Neon Transfection System (Thermo Fisher Scientific). Editing efficiency was determined as described previously (69, 74) using forward primer 5’ - GATACAGGGCTGGCTCTATGC-3’ and reverse primer 5’ - CAAGAGAGCCTTCCGAAAGAGG-3’.

### Doxycycline-inducible Cas9 treatments

Cas9 expression in the doxycycline-inducible system was validated by Western Blotting with an antibody against Cas9 (Cell Signaling Technology #14697S, 1:1000) and α-Tubulin loading control (Sigma #T9026, 1:10000). Cells were trypsinized, pelleted, washed with PBS, and lysed at 4 ºC in RIPA Buffer (Boston Bioproducts) supplemented with cOmplete mini protease inhibitor (Millipore Sigma), PhosSTOP protease inhibitor (Roche), 1 mM DTT, and 1 mM PMSF. Samples were centrifuged at 17,000 × g for 30 min at 4 ºC and the supernatant was run on a 10% Mini-PROTEAN TGX precast polyacrylamide gel (BioRad). Protein was transferred to a PVDF membrane using the iBlot 2 (Life Technologies). The membrane was blocked with 5% milk in TBST for 1 h at room temperature, followed by incubation with primary antibodies overnight at 4 ºC. Three washes were performed with TBST followed by 1 h incubation with secondary antibody (ECL, HRP linked, GE Healthcare) and another series of washes. Membranes were imaged using an ImageQuant LAS 4000 (GE Healthcare).

sgRNAs (Supplementary Table S1) were cloned into pLenti-Guide-Puro (Addgene) and delivered to hTERT-immortalized RPE-1 cells carrying a tetracycline-inducible promoter by lentiviral transduction, as above. Starting 24 h after transduction the population of cells was selected for one week in 12 µg/ml puromycin. For all experiments using the doxycycline-inducible system, cells were treated with 40 nM ON-TARGETplus siRNA SMARTpool L-003329-00-0050 (Dharmacon) to deplete p53 and prevent cell cycle arrest caused by Cas9 treatment or micronucleation. siRNA was transfected using Lipofectamine 3000 (Life Technologies) according to manufacturer’s instructions. 6 h after siRNA treatment cells were synchronized by serum starvation in 0.1% FBS-containing media. 24 h later, cells were released from this block into complete medium containing 0.5 µg/ml doxycycline, which was washed out 15 h later by 5 washes. When MPS1 inhibitor (1 µM NMS-P715, EMD Millipore) was used to produce micronuclei from mitotic errors, cells were instead released without doxycycline, and MPS1 inhibitor was added ∼18 h after release, before the next cell division. MPS1 inhibitor was then washed out by 5 washes with complete medium 20 h later. Cells were then transferred to coverslips or dishes for immunofluorescence or FISH experiments, live-imaging experiments, or plated for Look-Seq.

### Measurement of editing efficiency

DNA was isolated 48 h after RNP transfection or doxycycline washout using the PureLink Genomic DNA kit (Invitrogen) according to manufacturer’s instructions. PCR (50 µl reactions) was performed using Q5 High-Fidelity DNA Polymerase for 40 cycles after an initial 30 s denaturation step at 98 ºC [5 s 98 ºC, 10 s 60 ºC, 15 s 72 ºC] and a 2 min final extension at 72 ºC with 2.5 mM dNTP, 10 µM forward and reverse primers, 10 µL Q5 Reaction Buffer, and at least 20 ng of genomic DNA. 2% agarose gels were run in TAE buffer on an aliquot of PCR product to ensure production of a unique PCR product of the appropriate size (211–363 bp). Primer pairs utilized were as follows: Chr2p (5’ -CAAACGGCCACCGATGGAGAGGTCT- 3’ ; 5’ - CCCAGGTGTGCATAAGTAAGAGCAG-3’); Chr4q (5’ -GTGTATATAGTATATATAAATGGGC- 3’ ; 5’ -CCCCTTCCTACCTCTATCAACACAG-3’); Chr5q (5’ -GCTTCAGCAATCCTCTCGTC-3’ ; 5’ -CATATCACCCATCCCCTTTG-3’); Chr6q (5’ -GAAGTAGGGCATTTTTCTGATG-3’ ; 5’ - GAAGATTGATAAGCCATTTTGG-3’); ChrXq (5’ -TGTCATTTGCACTTGCTGAATCCAC-3’ ; 5’ - AGCATAGGTAAGGTAGTGACAAATA-3’). PCR products were purified using the QIAquick PCR purification kit (QIAGEN) and diluted to 20 ng/µL, as measured by Qubit dsDNA HS Assay Kit (Invitrogen). Samples were then submitted to Genewiz for Amplicon-EZ sequencing or to the Center for Computational and Integrative Biology DNA core facility of Massachusetts General Hospital for amplicon next generation sequencing. Analysis of the raw data for detecting CRISPR variants from NGS reads was performed with the algorithms from Genewiz Amplicon-EZ service or the MGH core and meta-analysis to estimate the percentage of editing efficiency was performed manually by the users.

### Fluorescence in situ hybridization of immortalized cell lines

FISH probes utilized in this study were as follows: Chr2p Subtelomere (Cytocell, LPT 02PG-A); Chr2 Centromere (Cytocell, LPE 002R-A); Chr4q Subtelomere (Cytocell, LPT 04QG-A); Chr4 Centromere (Cytocell, LPE 004R- A); Chr5q Subtelomere (Cytocell, LPT 05QG/R-A); Chr5 Centromere (Cytocell, LPE 005R-A); Chr6q Subtelomere (Cytocell LPT 06QR-A); ChrXq Subtelomere (Cytocell, LPT XYQG-A); ChrXq Centromere (Cytocell LPE 0XR-A)

Cells were seeded on #1.5 glass coverslips and were transfected with Cas9 RNP, as described. Cells were fixed ∼30 hours after release from starvation media, in the first interphase where the cells have formed micronuclei. Prior to fixation the coverslips were swelled in pre-warmed 75 mM KCl and incubated at 37 ºC for 20 min. Fixation was performed by dropwise addition of 0.5 volume of -20 ºC Carnoy’s solution (3:1 methanol:acetic acid). After 5 minutes, the solution was exchanged for fresh -20 ºC Carnoy’s solution twice more. Coverslips were then air dried for 48 h. Coverslips were warmed in 2X SSC + 0.5% NP-40 at 37 ºC for 30 min, and then dehydrated in ice cold solutions of 70, 85, and 100% ethanol for 2 minutes each. Subtelomerespecifc or centromeric probes were diluted 1:5–1:10 in hybridization buffer B (Cytocell, purchased by Rainbow Scientific # HB1000L) and applied to the samples after air drying. Coverslips were then sealed onto glass slides with rubber cement, denatured at 73 ºC for 2 min, and hybridized in a humidified chamber at 37 ºC for two days. After hybridization, coverslips were floated from the slides in a PBD solution composed of 0.1 M Na2HPO4, 0.1 M NaH2PO4, and 0.1% NP-40 for 3 min at RT. Samples were washed in 72 ºC 0.5 × 2X SSC + 0.5% NP-40 for 5 min, and then transferred to 2.5 µg/ml Hoechst 33342 (Life Technologies) dissolved in PBD solution for 10 min. Coverslips were then air dried and mounted on clean glass coverslips using ProLong Gold antifade (Life Technologies) or Vectashield Antifade Mounting Medium with DAPI (H-1200, Vector laboratories). Denaturation and wash steps were performed using a HybEZ II Hybridization system (ACD). Samples were imaged by confocal microscopy, as described below.

### Fluorescence in situ hybridization of CD34+ HSPCs

For detection of chr2 abnormalities, two BAC clones were used as probes, one located distal to the *BCL11A* locus (2p21) as the telomeric marker and a clone from 2q11.2 as the centromeric marker. The telomeric BAC DNA (hg19 chr2:47612794 - 47782780) was labeled with a red-dUTP (AF594, Molecular Probes) by nick translation and the centromeric BAC DNA (hg19 chr2:99969552 - 100200667) was labeled with a green-dUTP (AF488, Molecular Probes). Both labeled probes were combined with sheared human DNA and hybridized in a solution containing 50% formamide, 10% dextran sulfate, and 2X SSC. The cells were then stained with 4,6-diamidino-2-phenylindole (DAPI) and imaged using a Nikon Eclipse 80i with a 100 ×/1.40 NA Plan Apo objective and Cytovision version 7.7 (Leica Biosystems).

### SKY of CD34+ HSPCs

Day 1 post-electroporation, CD34+ cells were harvested by routine cytogenetic methods after a 4 h colcemid incubation. Commercially prepared SKY probes and protocols from Applied Spectral Imaging (Carlsbad, CA) were used for the hybridization and detection steps. Mitotic spreads were analyzed and breaks of chr2p were quantified.

### Indirect immunofluorescence microscopy

Immunofluorescence was performed as described with minor modifications (31, 44). Cells were seeded on #1.5 glass coverslips. For experiments measuring DNA replication by EdU incorporation, 10 µM EdU was added to complete medium 5 h before fixation. Cells were washed once with PBS and then fixed with 4% paraformaldehyde for 20 min. Fixed samples were washed 3 times with PBS, and permeabilized by PBS + 0.5% Triton X-100 for 5–10 min at room temperature and washed again with PBS. Samples were then blocked in PBS + 3% BSA for one hour. EdU staining was performed following instructions from the Click-iT Plus EdU Cell Proliferation Kit for Imaging (Invitrogen). Samples were incubated with primary antibody for one hour, washed three times with PBS + 0.05% Triton X-100, and incubated with secondary antibody for 45 min, followed by another series of 3 washes in PBS + 0.05% Triton X-100. Next, samples were incubated in PBS + 2.5 µg/ml Hoechst 33342 (Life Technologies) for 10 minutes, followed by two washes in PBS. Finally coverslips were mounted on glass slides using ProLong Gold antifade (Life Technologies). Primary antibodies: γH2AX (1:400-500, MilliporeSigma, 05-636-I, LBR (1:100, Abcam, ab32535). Secondary antibodies: Alexa Fluor, 488, 568 and 647 (1:1000, Life Technologies).

Confocal images were collected using a Nikon Ti-E inverted microscope with a Yokogawa CSU-22 spinning disk head with the Borealis modification. Z-stacks were collected for 9 images at 0.4-0.6 µm spacing using a CoolSnap HQ2 CCD camera (Photometrics) and a 60 ×/1.40 NA Plan Apo oil immersion objective (Nikon). Alternatively, a Ti2 inverted microscope fitted with a CSU-W1 spinning disk system (Nikon) was used. Z-stacks were collected to cover the whole volume of cells at 0.4-0.6 µm spacing using a Zyla 4.2 sCMOS camera (Andor) and a 60 ×/1.40 NA Plan Apo λ oil objective.

### Indirect immunofluorescence microscopy of CD34+ HSPCs

1.5 × 105 CD34+ cells were deposited on glass slides by centrifugation using a Cytospin(tm) 4 cytocentrifuge (Thermo Scientific) for 5 min at 800 rpm. Fixation and indirect immunofluorescence were performed as above. Images were acquired with single-plane widefield illumination on a Nikon Eclipse Ni microscope using Nikon NIS-Elements software and a 40 ×/0.75 Plan Fluor objective. Antibodies were the same as those listed above.

### HbF Quantification

Fetal hemoglobin quantification by ion-exchange high-performance liquid chromatography was performed as previously described (69). Erythroid differentiated CD34+ cells were lysed with hemolysate reagent (Helena Laboratories) and analyzed using ion-exchange columns on a Prominence HPLC System (Shimadzu Corporation). Proteins eluted from the column were identified at 220 nm and 415 nm with a diode array detector. The relative amounts were calculated from the area under the 415 nm peak and normalized based on the DMSO control. Hemoglobin peaks were identified using commercially available standards (Helena Laboratories).

### Live-cell imaging

Live-cell imaging was performed as described previously with minor modifications (31, 44). For live-cell imaging experiments to investigate bridge formation, doxycycline-inducible Cas9 expressing RPE-1 cells expressing eGFP-BAF and the guide targeting chr5q were plated on ibiTreat 24-well µ-Plates (ibidi) and placed on a Nikon inverted microscope (Ti-E or Ti2) with Perfect Focus for widefield microscopy. The microscope was equipped with an environmental chamber to maintain cells at 37 ºC and with humidified 5% CO2. Imaging was performed in 20 min intervals using a 20 ×/0.75 NA Plan Apochromat Lambda objective (Nikon) and Z-stacks of three images with a 2 µm step size. Images were acquired using a Zyla 4.2 sCMOS camera (Andor) for up to ∼50 h.

### Image analysis

NIS-Elements (Nikon) was used to analyze live-cell imaging videos, and ImageJ was used to create annotated videos. Quantitative image analysis for fixed-cell experiments was performed using ImageJ. Briefly, nuclear segmentation was performed on maximum intensity projections based on Hoechst staining. This segmentation was used as a mask, and, if necessary, the mask was manually refined by the “Watershed”, “Erode”, or “Draw” functions. These masks were then applied to maximum intensity projections of other channels to measure the mean fluorescence intensity of channel. Background subtraction was performed by measuring the mean fluorescence intensity of a square region near the primary nucleus and micronucleus. Analysis of micronucleus formation and DNA damage in CD34+ HSPCs was performed qualitatively by sample-blinded individuals for the presence or absence of a single large focus of γH2AX signal covering most of the micronucleus.

### Live-cell imaging followed by single cell isolation and single cell whole genome sequencing

Look-Seq was performed as previously described (30, 31). Briefly, single cells were seeded by flow sorting in 384-well µClear plates (Greiner), or were seeded in bulk on a 35-mm gridded ibiTreat dish (ibidi) and eGFP-H2B and TDRFP-NLS were imaged by widefield fluorescence imaging in intervals of 10–15 min, as above. After allowing sufficient time for micronucleus-containing cells to divide daughter cells of interest were separated (∼40 h after micronucleus formation). Cells were considered to have reincorporated their micronuclei if no fragments of GFP-H2B were detected in the cytoplasm after division. For cells in 384-well plates, separation was performed by trypsinization followed by limited dilution of daughters into a new 384-well plate (31). For cells on ibiTreat dishes, the dish was transferred to another Nikon inverted microscope that was equipped with a CellEctor single-cell isolation system (Molecular Machines and Industries) and cell adhesion was loosened by exchanging the culture medium with a PBS-based non-enzymatic dissociation reagent (Sigma). Within ∼30 min of applying dissociation reagent, cells of interest from live-cell imaging were identified and directly picked from the imaging dish using a robotically controlled glass capillary with an inner diameter of 40 µm that aspirated 80 nL of volume (Molecular Machines & Industries). This volume containing the cell was then deposited into a 5 µL droplet of PBS contained in a PCR-tube cap. Whole genome amplification was performed using the REPLI-g Single Cell kit (QIAGEN), with initial lysis steps being performed in the PCR-tube lids, and amplification terminated after 80 min. Amplified DNA was then purified and sheared by sonication (Covaris) into ∼500 bp fragments. Sheared DNA was then processed by a Library Preparation Kit (KAPA) to prepare for multiplexed next-generation sequencing as previously described (31).

### Quality assessment of sequencing libraries

Library quality assessment was performed as described previously (30, 31). Briefly, before deep-sequencing libraries were subjected to low-pass sequencing (∼0.1 × genome coverage) by the MiSeq platform (Illumina). From this we visually assessed library quality by the uniformity of whole-genome amplification in 10 Mb bins. Furthermore, low-pass sequencing was used to assess haplotype-specific DNA copy number in order to identify cells with missegregation of the targeted chromosome. Libraries that passed quality checks were then subjected to deep sequencing (8-47 × genome coverage; 19 × mean coverage, 11 × median coverage) by HiSeq 2500 (Illumina) or NovaSeq 6000 platforms (Illumina).

### Sequencing data processing and haplotype-specific DNA copy-number calculation

Sequencing data processing and haplotype-specific DNA copy-number analysis were carried out using the same bioinformatics pipeline and computational workflow as described previously (30, 31).

### Structural variant (SV) detection in single-cell genomes

Structural variants were detected using our previously described pipeline (30, 31). As the Phi29 polymerase produces artificial chimeras between loci within close proximity due to template switching, we excluded short-range intra-chromosomal discordant reads (distance between fragments < 10 kb). Because of this limitation, we do not detect short-range SV events, including insertion/deletion events at the CRISPR cut sites and local fold-back events that are expected to be generated between fused sister chromatids. We previously reported that false structural variants due to chimeric DNA generated by single-cell whole-genome amplification are enriched between loci separated by ≤ 150 kb (31). By contrast, most de novo rearrangements resulting from DNA damage from micronuclei are formed between loci separated by ≥ 1 Mb. To exclude false SV events due to artificial chimeras with a stringent threshold, we only considered long-range intrachromosomal SVs with breakpoints separated by ≥ 1 Mb in this study.

In addition to the above described short-range chimeric DNA fragments occurring within amplicons, whole-genome amplification also generates random chimeric fragments between amplified DNA. As such chimeras are generated between random amplified DNA rather than from the original DNA template, the allele fraction of random chimeric fragments should be lower than the allele fraction of reads supporting true structural variants. Due to variation in the sequence coverage, we filtered low allele fraction variants using sample-specific read-depth cutoffs determined from the average allelic coverage. The average allelic coverage at different read-depth cutoffs was determined from the allelic depths at heterozygous sites in the parental RPE-1 line (31). In each sample, we determined the allelic depth at which the average allelic coverage (detection sensitivity for a genetic variant on a single chromosome) is approximately 50% and used this value as the minimum read-depth cutoff if it is above three. The requirement of at least three supporting reads is necessary for excluding random chimeras generated during library construction. For interchromosomal events, we required at least one of the three supporting reads to be a split alignment and one to be a discordant read pair mapping. We note that the conclusions regarding statistical enrichment of structural variants within each sample are not dependent on the read-depth cutoff, but the choice of sample-specific readdepth cutoffs allows consistent estimation and comparison of the frequency of DNA breaks across different samples with varying uniformity and sequencing depth.

### Poisson test for breakpoint enrichment, and definition of fragmentation

We performed two-sample onesided Poisson tests to determine whether SVs are enriched on the CRISPR-targeted segment compared to the background rate of SVs across the genome. We calculated this statistic relative to the depth of sequencing coverage in the targeted region, as follows. For each pair of granddaughter cells (*a, b*), we calculated the fraction of reads (*r*_*a*_, *r*_*b*_) mapping to the genomic interval telomeric to the CRISPR cut in each sample. The null hypothesis is that breakpoints are drawn according to a Poisson process with Poisson parameter *λ* = total number of breakpoints observed in the pair. We computed the probability of the observed number of breakpoints on the targeted segment occurring equal to *t* = (*r*_*a*_ + *r*_*b*_)*/*2, conditioned on the total number of breakpoints. The test is implemented as a one-sided, one-sample binomial test *P* (*X* ≥ *k*) where *n* = total breakpoints observed across the pair, *P* = (*r*_*a*_ + *r*_*b*_)*/*2, and *k* = breakpoints observed on the targeted segment.

Segments were considered fragmented by visual inspection of copy number plots for the presence stretches of allelespecific reciprocal copy number change between daughter cells or many rearrangements on the targeted arm in both daughters. For Fig. 2, individual cells were marked as having ‘clustered rearrangements’ if there was significant enrichment by the Poisson test, the daughter did not lose the missegregated allele, and there was at least one rearrangement found in the cell in cases of fragmentation.

## Supplementary Figures

**Table S1.**
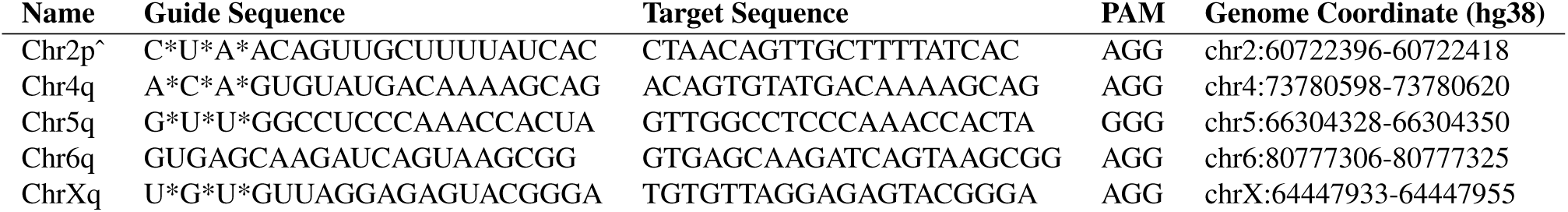
Summary of gRNAs. Table of gRNA sequences and coordinates used in this study. * For RNP treatments, modified 2 -O-Methyl bases with phosphorothioate linkages. ^ From Wu et al., 2019

**Fig. S1.**
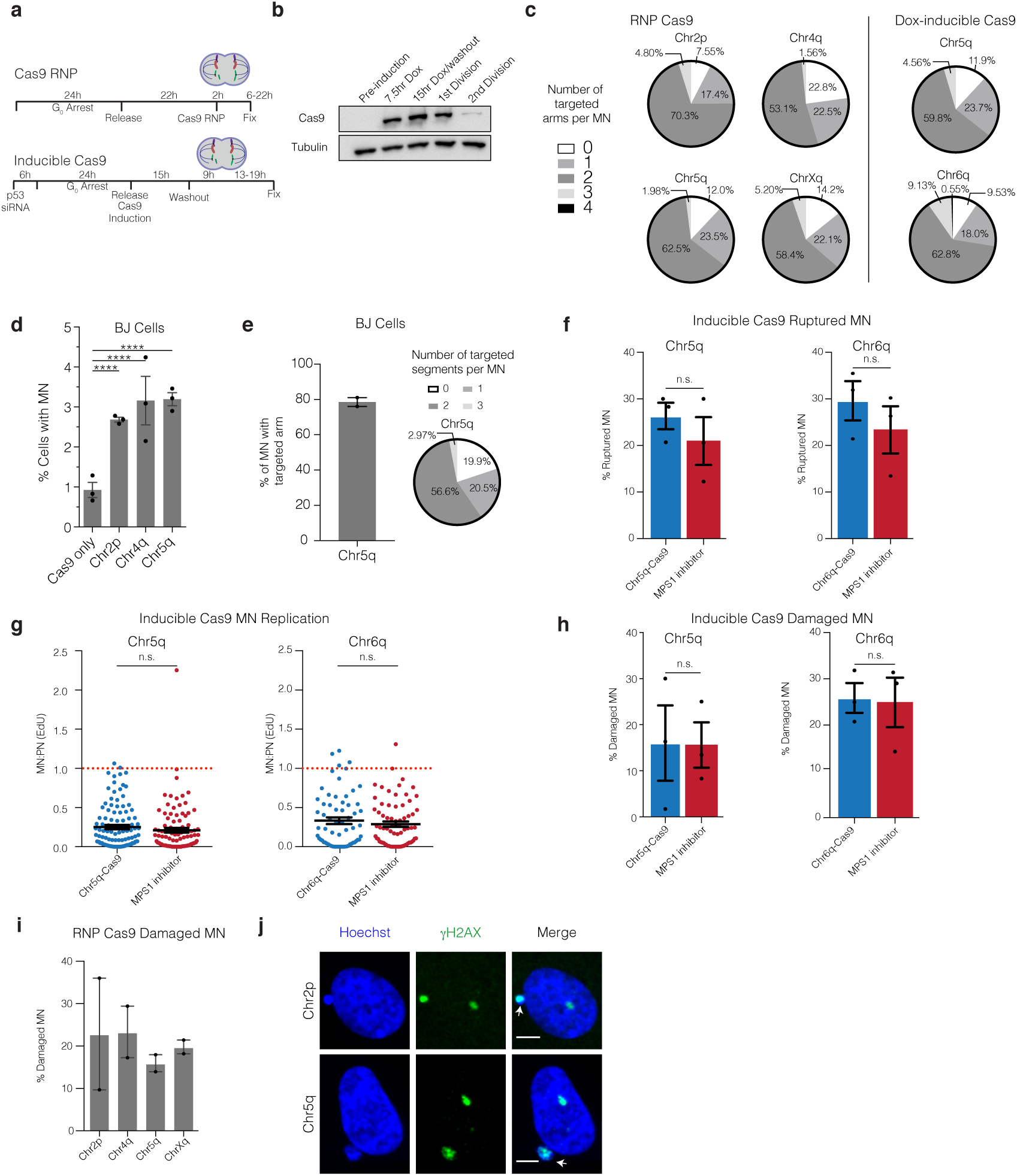
Micronucleus formation and DNA damage after CRISPR-Cas9 genome editing in several cell lines. (a) Scheme of the RNP transfection and inducible Cas9 expression genome-editing experiments. Top, RNP transfection experimental scheme. Bottom, inducible Cas9 expression with constitutive expression of gRNAs (RPE-1 cells) experimental scheme; G0 cell cycle block was by serum starvation. Dividing cell cartoon represents approximate time of cell division. (b) Representative Western blot to detect Cas9 levels at the indicated intervals before and after induction. 1st division is 24 hours after serum starve release, and 2nd division is 48 hours after release. Dox is doxycycline. *n* = 3 experiments. (c) Number of cleaved chromosome arms contained within micronuclei for the indicated gRNAs and Cas9 expression strategies (RPE-1 cells). Arm number was determined by FISH to detect the centromere (RNP Cas9) and/or subtelomere of the targeted chromosome (RNP Cas9 and Dox-inducible Cas9). RNP Cas9: for 2p: *n* = 2 experiments with 64 micronuclei counted, 4q: *n* = 2 experiments with 58 micronuclei counted, 5q: n= 3 experiments with116 micronuclei counted, Xq: *n* = 2 experiments with 96 micronuclei counted; (Dox) Doxycycline-inducible Cas9; *n* = 3 experiments; 186 micronuclei counted per condition. (d) Frequency of micronucleation in synchronized BJ fibroblasts after RNP transfection; (*n* = 3 experiments with 2378, 2487, 2423, 2714 cells, left to right). Error bars: mean *±* SEM, *P <* 0.0001, two-tailed Fisher’s exact test. (e) Left, percentage of MN containing the targeted chromosome arm for the chr5q-targeting gRNA, as counted using subtelomeric FISH probes. Right, pie chart of the number of chr5q chromosome arms per micronucleus in BJ cells, as determined from centromere-specific and subtelomere-specific FISH probes. (*n* = 2 experiments counting 109 micronuclei) Error bars: mean *±* SEM. (f) Nuclear envelope rupture frequency for CRISPR-MN as compared to spindle checkpoint inhibitor-induced micronuclei (MPS1 inhibition, primarily containing whole chromosomes). Rupture was identified by heavy accumulation of the inner nuclear envelope protein lamin B receptor (LBR)44, which is measured from the fluorescence intensity of LBR labeling in micronuclei relative to the intensity in primary nuclei. Rupture was operationally defined as an MN:PN ratio > 3 (*n* = 3 experiments with 201 and 167 micronuclei analyzed for chr5q, p = 0.2216 and 165 and 152 micronuclei counted for chr6q, p = 0.2034). Error bars: mean *±* SEM, two-tailed Fisher’s exact test. (g) DNA replication defect of CRISPR-MN. EdU fluorescence intensity was measured after a 5-hour pulse. Only cells that had entered S-phase were scored (operationally defined as > 150 a.u. EdU signal in primary nucleus). Dotted red line is normal levels of DNA replication in the micronucleus relative to the primary nucleus (*n* = 3 experiments with 109 and 97 micronucleated cells analyzed for chr5q, p = 0.1698 and 65 and 73 micronucleated cells analyzed for chr6q, *P* = 0.6948). Error bars: mean *±* SEM; two-tailed Mann-Whitney U-test. (h) CRISPR-MN acquire DNA damage. Shown is the frequency of γH2AX positive micronuclei (> 3 standard deviations above the average signal in primary nuclei) for the indicated gRNAs using the inducible Cas9 system (*n* = 3 experiments with 203 and 184 micronucleated cells analyzed for chr5q, *P* = 0.6870 and 175 and 169 cells analyzed for chr6q, *P* = 0.8053). Error bars: mean ± SEM, two-tailed Fisher’s exact test. (i) CRISPR-MN acquire DNA damage. Shown is the frequency of γH2AX positive micronuclei (> 3 standard deviations above the average signal in primary nuclei) for the indicated gRNAs using the RNP Cas9 system (*n* = 2 experiments with 56, 46, 82, and 50 micronucleated cells analyzed, left to right). Error bars: mean ± SEM. (j) Example images showing γH2AX labeling (RNP lipofected RPE-1 cells). White arrows: micronuclei. Scale bars, 5 µm. The γH2AX focus in the primary nucleus likely decorates the centric portion of the broken chromosome. Alternatively, or additionally, it may label a DNA break on the homologue.

**Fig. S2.**
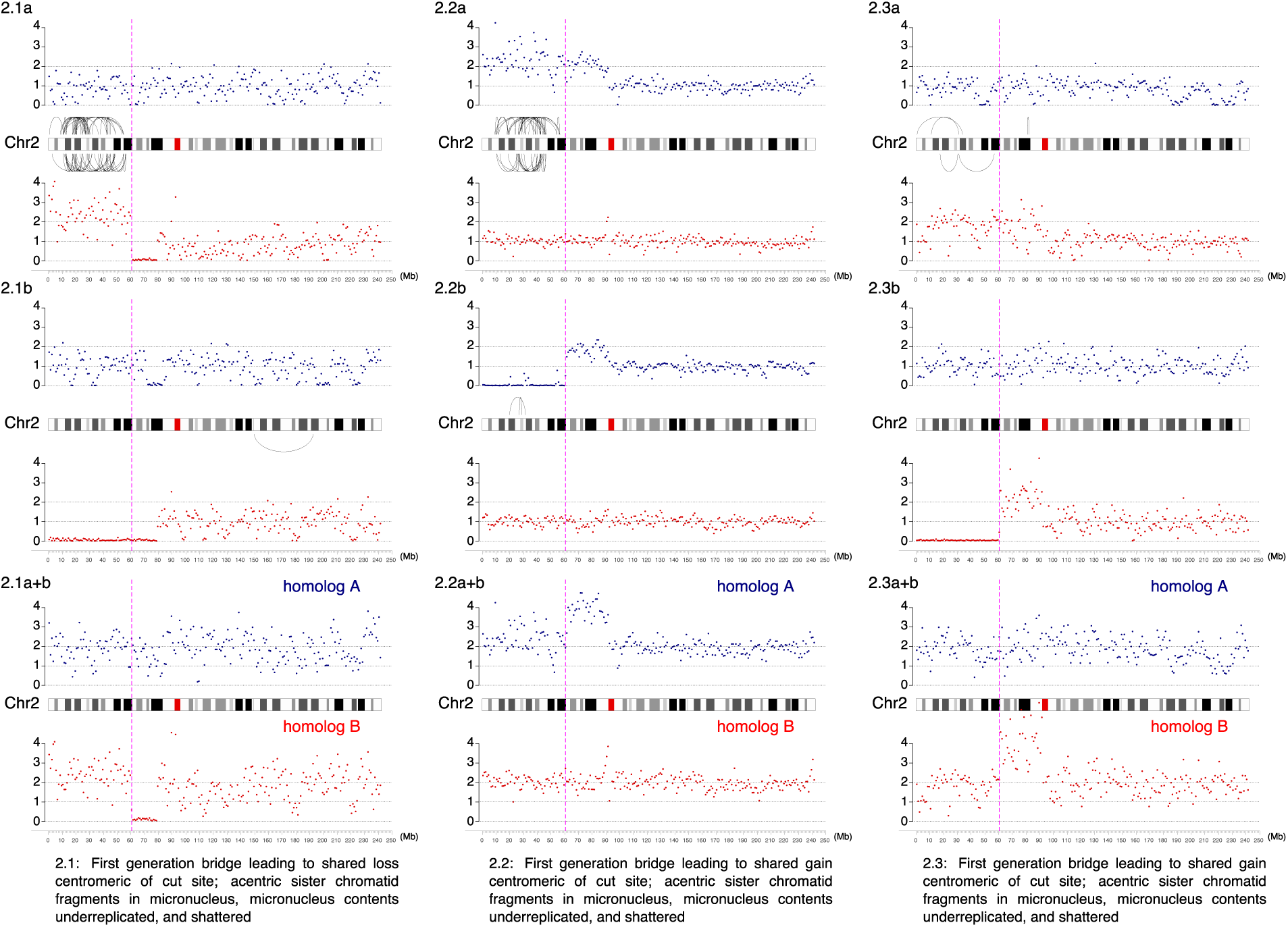
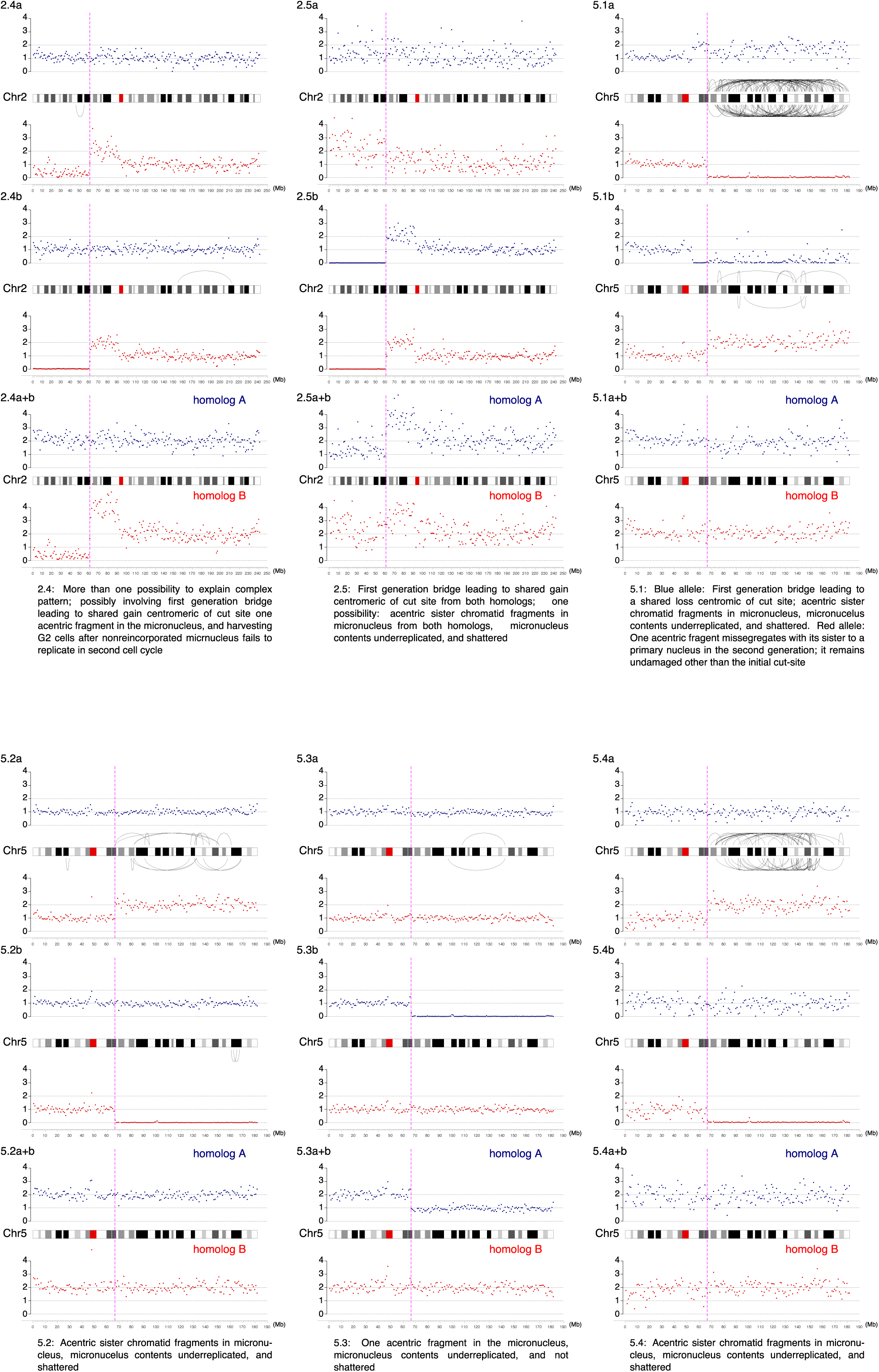
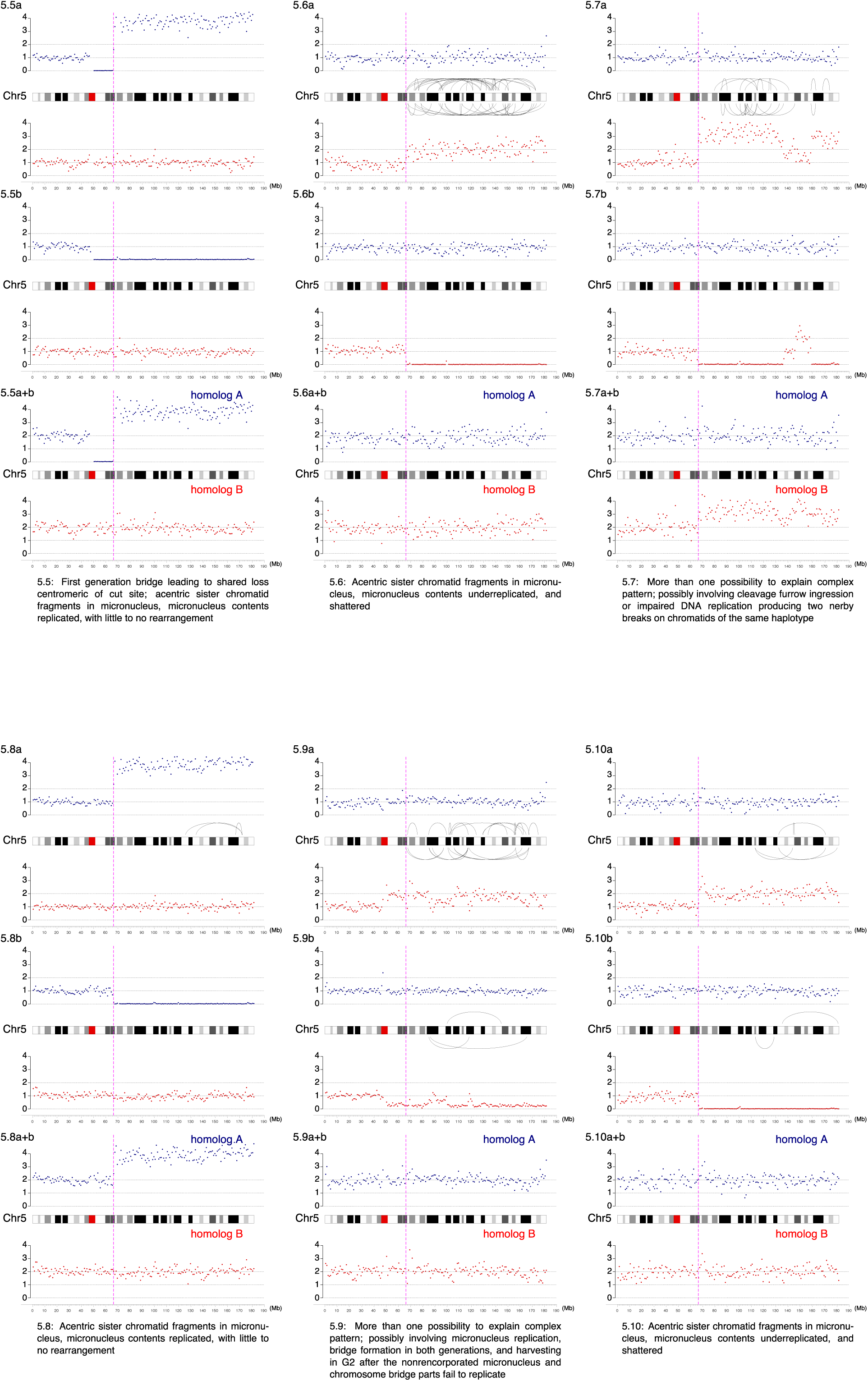
Haplotype copy number and SVs for the targeted chromosome for each sample in the paper. Haplotype-resolved copy number and structural variant analysis for the targeted chromosome for each granddaughter pair. Red and blue dots represent 1 Mb copy number bins for each homolog, and curved lines represent structural variants of *≥* 1 Mb that could be on either homolog. Top, ‘granddaughter a’; middle, ‘granddaughter b’; bottom, sum copy number for each homolog for the pair of cells. Note that in most cases there should be a total of two red and two blue copies per granddaughter pair, and deviation from this represents certain missegregation or events, such as first-generation bridge formation. Text: inferred most likely explanation for each copy number and rearrangement profile. Note that alternative explanations exist for many samples, such as a G1 cut followed by replication of the cut chromosome.

**Fig. S3.**
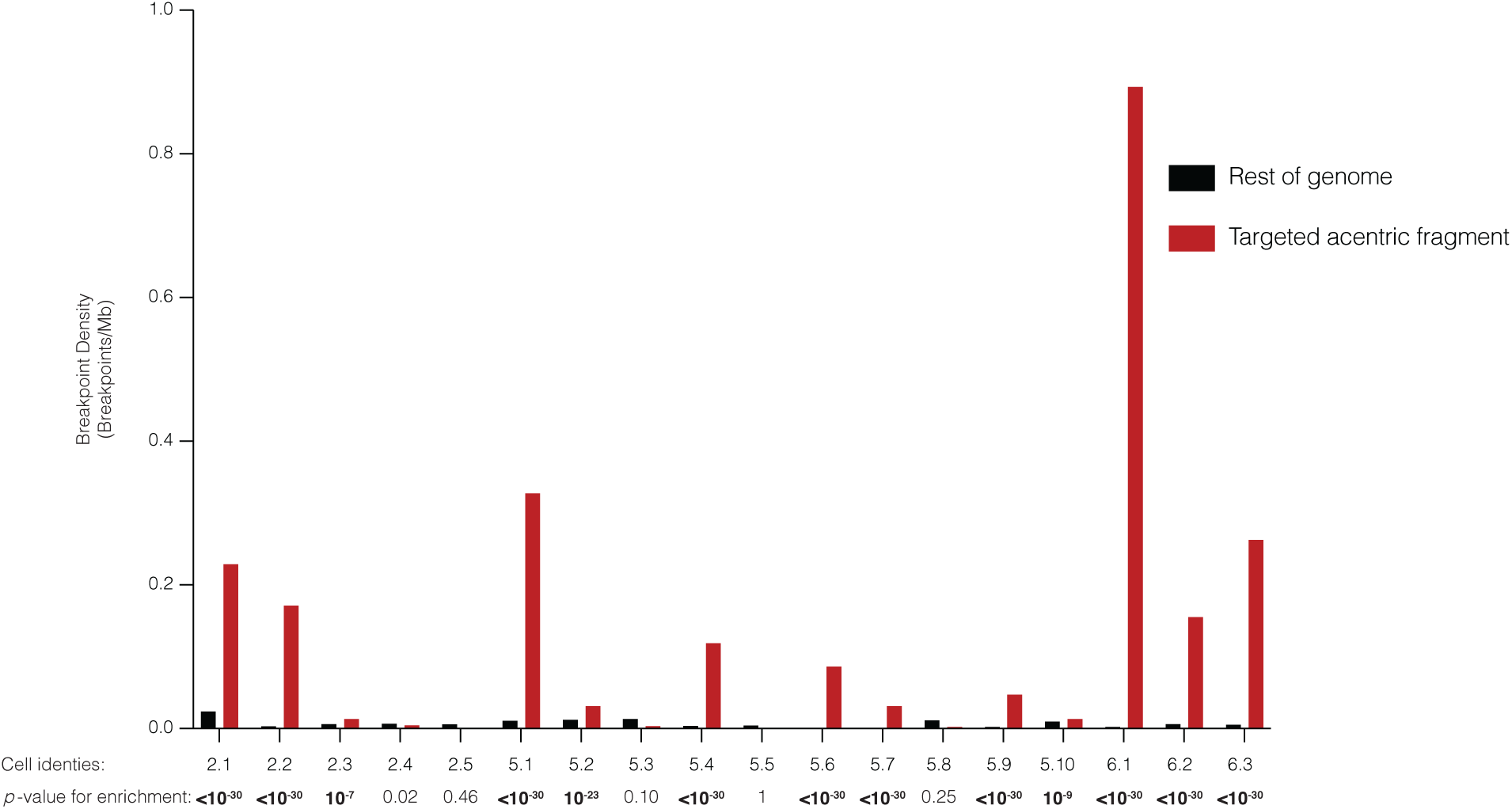
Clustering of DNA breakpoints, indicative of chromothripsis, to the telomeric side of the CRISPR-Cas9-targeted chromosome arm. Breakpoint density for each daughter pair telomeric of the cut-site (red), relative to the rest of the genome (gray), normalized by read depth. Significance is derived from a one-sided Poisson test (Zhang et al., 2015). *P* values are rounded to the nearest exponent, except for those *<* 10^*–*30^. Bolded *P* values denote significance after Bonferroni correction. Bonferroni-corrected *α* = 0.0028.

**Fig. S4.**
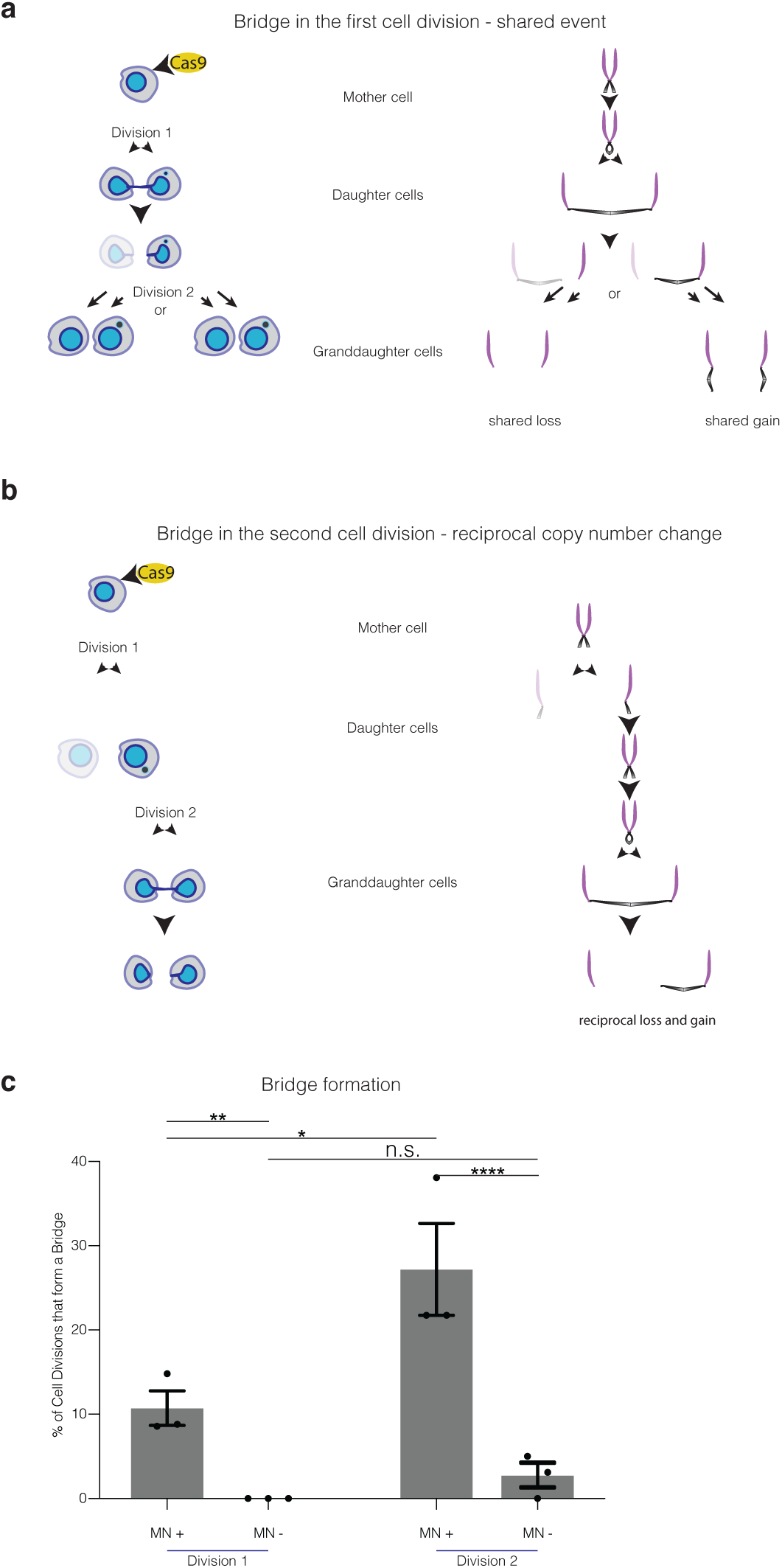
Chromosome bridge formation after CRISPR-Cas9 genome editing. a) A bridge formed during the first cell division after Cas9 addition will yield shared losses (left granddaughter pair) or gains (right granddaughter pair) depending upon how the bridge breaks. Note that this copy number alteration will be on the centromeric side of the CRISPR-Cas9 break. Cells and chromosomes are depicted as in Fig. 2. To focus on the fate of the progeny of the micronucleated daughter, the non-micronucleated daughter cell is faded and not followed. In this example, the micronuclear chromosome from the first division is not reincorporated becomes a micronucleus in one granddaughter. b) A bridge formed in the second cell division will yield reciprocal copy number gains and losses on the centromeric side of the break (comparing the granddaughters). To focus on the fate of the micronucleated daughter, the non-micronucleated daughter cell is faded and not followed. c) The frequency of detectable chromosome bridges by live-cell imaging after CRISPR-Cas9 genome editing in RPE-1 cells expressing a fluorescence reporter that marks chromosome bridges efficiently (GFP-BAF). DNA breaks were induced with the Chr5q-targeting inducible Cas9 system. Chromosome bridges frequently arise when a micronucleus forms in at least one daughter cell in the first division (MN+), whereas when a micronucleus is not formed, bridge formation is uncommon (MN-). In the second division, micronucleated cells are more prone to bridge formation (MN+) as compared to non-micronucleated cells (MN-). Additionally, bridge formation is more frequent in the second division, which may be explained by isolation of the acentric arm from the centric fragment of the chromosome (*n* = 3 experiments with 96 and 97 cell divisions imaged [division 1] and 67 and 76 divisions imaged [division 2]). Error bars: mean *±* SEM, ** *P* = 0.0015; **** *P <* 0.0001; * *P* = 0.0104, n.s. *P* = 0.1916, two-tailed Fisher’s exact test.

